# Cell lipotypes localization in brain by mass spectrometry imaging

**DOI:** 10.1101/2024.02.02.578599

**Authors:** Jonatan Martínez-Gardeazabal, Marta Moreno-Rodríguez, Alberto Llorente-Ovejero, Estibaliz Gonález de San Román, Laura Lombardero, Iker Bengoetxea de Tena, Juan Sustacha, Carlos Matute, Iván Manuel, Paolo Bonifazi, Rafael Rodríguez-Puertas

**Affiliations:** Department of Pharmacology, Faculty of Medicine and Nursing. University of the Basque Country (UPV/EHU), Leioa, Spain; Computational Neuroimaging Lab, BioBizkaia Health Research Institute, Barakaldo, Spain; Achucarro Basque Center for Neuroscience, Leioa, Spain; Department of Neuroscience, Faculty of Medicine and Nursing. University of the Basque Country (UPV/EHU), Leioa, Spain; Centro de Investigación Biomédica en Red sobre Enfermedades Neurodegenerativas (CIBERNED), Madrid, Spain; Neurodegenerative Diseases, BioBizkaia Health Research Institute, Barakaldo, Spain; IKERBASQUE: The Basque Foundation for Science, Bilbao, Spain

## Abstract

The study investigates brain lipid super-specialisation by defining characteristic spectral lipotypic profiles for the five primary cerebral cell-types. Utilizing a computational approach, the research visualizes the anatomical distribution of these profiles with high spatial resolution in brain tissues. This method unveils cellular stereotypic lipidic signatures within the CNS, providing a new framework for exploring the physiological roles of lipids in diverse cell-types present in brain or in any other tissue.

---

The brain can be considered primarily as a lipid organ, composed mainly of phospholipids, sphingolipids, and cholesterol, which may constitute 50-60% of cell membranes^1,2,3^. These lipid molecules have a wide range of functions, including structural roles in cell membranes where various membrane proteins (e.g., receptors and ion channels) are anchored, but they are also versatile in metabolic and bioenergetic processes within cells. In addition, membrane lipids have been identified as reservoirs of lipid mediators and signalling molecules that act as neuromodulators to coordinate the above-mentioned processes. The development of powerful analytical techniques during the last decade, in particular mass spectrometry, notably imaging, has allowed for the anatomically resolved identification of lipids. Furthermore, cell types have been described to have specific lipid compositions as measured by UHPLC^4^. As a result, a super-specialisation of lipids in the brain has been revealed^5,6^, which reaches its highest expression when considering the recent lipotype hypothesis, which explains that distinct cellular lipidomes are both a consequence and a component of differentiation programmes at the cellular level, providing insight into the role of cellular lipid composition in differentiation and development, or even linking the lipotype to the ultimate cell fate^7^.

So far, the localisation of different cell types in tissues has been achieved by labelling specific proteins of each cell type using techniques such as immunohistochemistry or, more recently, based on their transcriptome e.g. MERFISH^8^. In our study, we focused on developing a method to anatomically localise cell types based on their lipotypic spectral signature. This approach has allowed us to determine their distribution without the use of other cell-specific markers and considering the information provided by the full lipid spectra rather than relying on single lipid-markers or proteins (as in standard immunochemistry).

In this study, we used purified rat cell-type samples obtained from cultures representing five representative major cell types present in the brain, specifically neurons (NEU), astrocytes (AS), oligodendrocytes (OLI), microglia (MG), and choroid plexus cells (PLE) (Fig. 1a). This work aimed to provide a proof-of-principle that, given the spectral lipid profile of selected cell types, it is possible to reconstruct their anatomical localisation by quantifying their contribution on local mass spectrum acquired from brain tissue. Therefore, we performed spectral lipotypic analysis (Fig. 1a) using matrix-assisted laser desorption/ionisation-assisted mass spectrometry imaging (MALDI-MSI) on the different samples investigated, specifically rat purified cell-types, human hippocampus, mouse and rat brain slices in healthy condition or in toxin-induced lesion (see methods). We first obtained high spatial resolution images, where each pixel contained the lipotypic spectral information of the different cell-types deposited on glass slides, and then used these data to reconstruct anatomical images of cell-type composition in tissue sections (Fig. 1a).

**Fig. 1.**
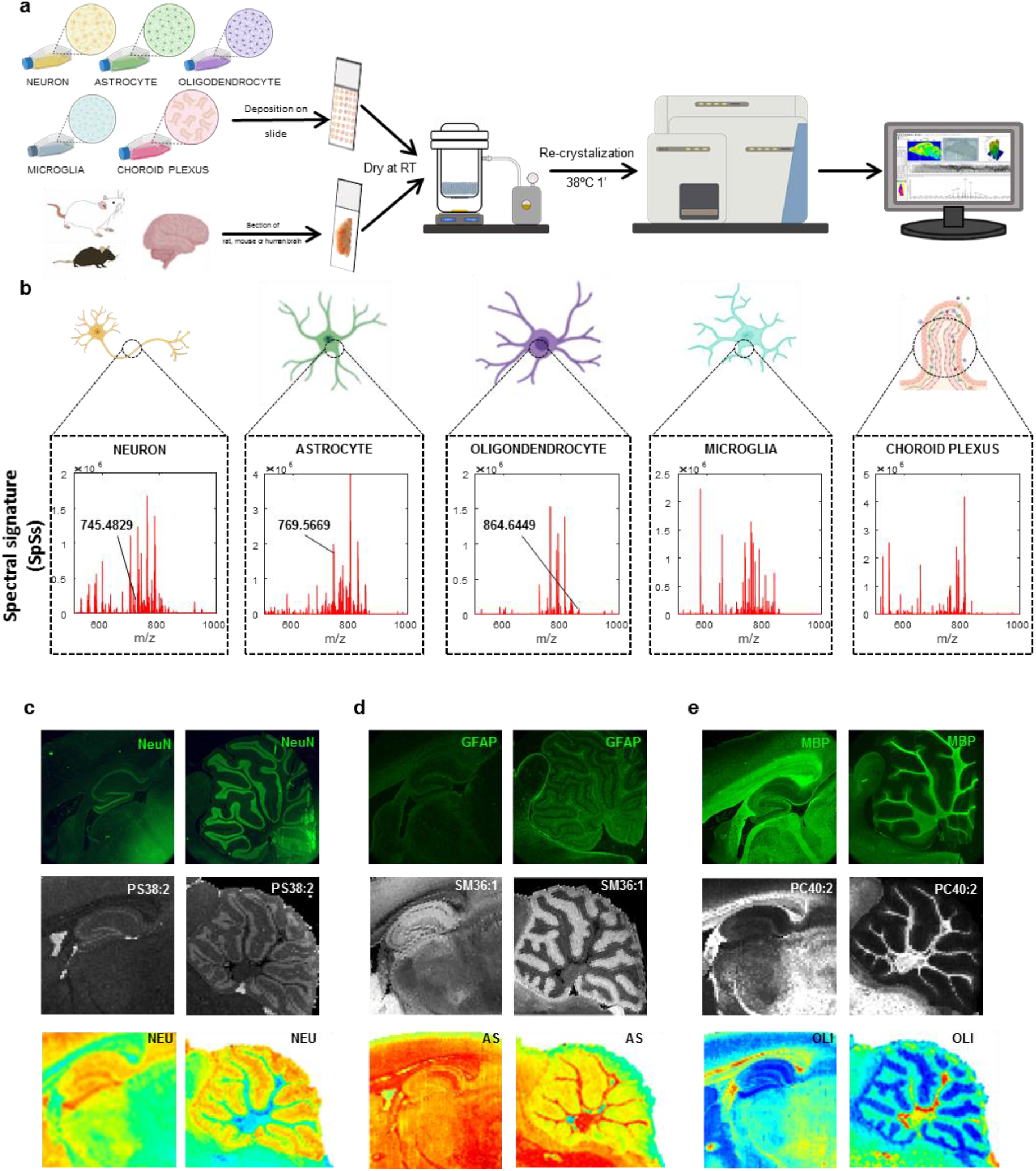
**a**, Workflow for sample processing by matrix-assisted laser desorption/ionisation-assisted mass spectrometry imaging (MALDI-MSI). **b**, Graphical representation of the lipotypes spectrum of the different cell types. **c**, NeuN immunoreactive neurons (top row) in hippocampus (left column, magnification x250) and cerebellum (right column, magnification x250), second row shows the distribution of the lipid PS(38:2)+H-H_2_O^+^ (m/z 745.4829), and bottom row shows reconstructed image from neuron lipotype, note that this image shows a wider distribution than the NeuN immunolabeling that is restricted to the nucleus or soma of the neurons. **d**, GFAP immunoreactive astrocytes (top row) in hippocampus (left column) and cerebellum (right column), second row shows distribution of lipid SM(d36:1)+K^+^ (m/z 769.5669) and bottom row shows reconstructed image from astrocyte lipotype. **e**, MBP immunoreactive (top row) oligodendrocytes in hippocampus (left column) and cerebellum (right column), second row shows distribution of lipid PC(40:2)+Na^+^ (m/z 864.6449) and bottom row shows reconstructed image from oligodendrocyte lipotype.

As first step, we analysed the MALDI-MSI results to extract specific information regarding the lipotypes of the five cell-types. Given a cell type (e.g. microglia), this process involved first image segmentation to define regions with sufficient density of cells (Extended Data Fig. 1a), displaying maximum power of the spectral signals associated to lipid signatures compared to background matrix (Extended Data Fig. 1b). Next, average spectral representation (AvSs) for a given cell type was computed across cultures (top blue spectrums in Extended Data Fig. 2). The final spectral signatures (SpSs) for each cell type (red spectrums in Fig. 1b and in Extended Data Fig. 2a) were obtained by removing linearly spectral contributions from other cell-types (see Methods and Extended Data Fig. 2). Note that while original AvSs (blue spectrums in Extended Data Fig. 2a) displayed in average high similarity (0.56 +/-0.09 SEM; see super-diagonal part of the matrix shown in Extended Data Fig. 2c and corresponding blue scatterplots between the cell-types spectra in Extended Data Fig. 2b; cosine similarity was used, see Methods), the similarity between the SpSs dropped down to 0.14+/-0.02 SEM (see sub-diagonals of Extended Data Fig. 2b,c), while preserving for each cell type high similarity to its original AvSs (0.79+/-0.05 SEM; see diagonals of Extended Data Fig. 2b,c). Overall, the above analysis shows that SpSs are characteristics for each cell type with low similarity between cell-types, while preserving the information of the original AvSs.

We then examined the similarity between the anatomical distribution of representative single lipids (Fig. 1c-e middle images), which were selected from the unique masses of each cell type, and the anatomical distribution of antibodies labelling proteins specifically expressed in the same cell types (Fig. 1c-e top images). The similarity, shown in Fig. 1c-e for a rat brain slice, was particularly evident in the anatomical distribution of lipids, associated with neurons labelled with antibodies to neuronal nuclear marker (NeuN) (Fig. 1c), astrocytes labelled with antibodies to glial fibrillary acidic protein (GFAP) (Fig. 1d) and oligodendrocytes labelled with antibodies to myelin basic protein (MBP) (Fig. 1e).

Next, in the same brain regions we used the SpSs, i.e. the characteristic spectral lipotypes for the five different cell-types, which provide a broader information on the cellular lipidic presence, to reconstruct anatomical images of cell-type distributions (bottom). To this end, we applied a general linear model based on the SpSs to quantify the abundance of a given cell type with spatial resolution of the scanned image. The results for comparison are shown on the bottom of Fig. 1c-e, confirming that the SpSs of each cell type allowed their differentiation in the subsequently reconstructed image.

In particular, the SpSs methodology allowed us to reconstruct an image of a control rat tissue section (Fig. 2a-f) for each cell type, providing information on their anatomical distribution without any molecular labelling. Furthermore, this analytical approach was applied under conditions of injury, allowing the visualisation of modifications in the cellular localisation of lipotypes in specific brain areas (Extended Data Fig. 3). We demonstrated in previous studies using this type of lesion increased Iba-1 and decreased GFAP labelling at the toxin injection site^9^. Lipotype-based reconstruction imaging gave results comparable to immunofluorescence techniques (Extended Data Fig. 3h,j). In addition, SpSs analysis of choroid plexus cells showed an increase in these cell types at the site of toxin delivery (Extended Data Fig. 3k). It is noteworthy that the information obtained from the spectral lipotypes of cultured rat cells can be used to reconstruct localisation in mouse brain slices (Fig. 2g-l). Furthermore, this algorithm has also been applied to human brain tissue, specifically in the hippocampus, revealing reconstructed image distributions that resemble those corresponding to the different cell types (Fig. 2m-r).

**Fig. 2.**
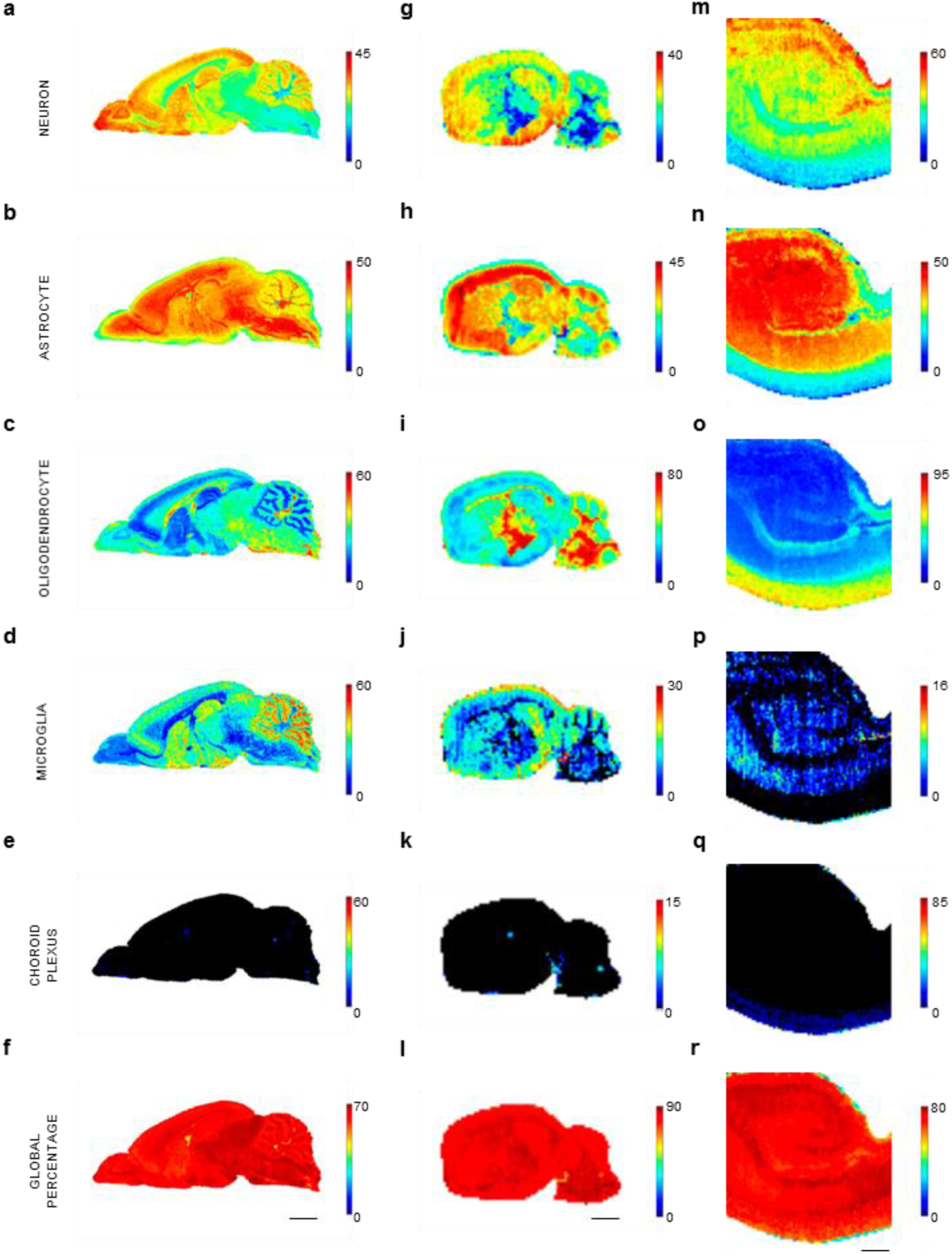
Reconstructed images of the localisation of the different lipotypes in rat (right), mouse (center) and human hippocampus (left) brain slices. **a-f**, Control rat brain in sagittal section, scale bar=4mm. **g-l**, Sagittal control mouse brain, scale bar=3mm. **m-r**, Human hippocampus, scale bar=5mm. **a**,**g**,**m**, Image reconstructed from the neuron lipotype. **b**,**h**,**n**, Image reconstructed from the astrocyte lipotype. **c**,**i**,**o**, Image reconstructed from the oligodendrocyte lipotype. **d**,**j**,**p**, Image reconstructed from the microglia lipotype. **e**,**k**,**q**, Image reconstructed from the choroid plexus lipotype. **f**,**l**,**r**, Reconstructed image illustrating the global percentage that could be explained as a proportion attributed in each pixel based on all the lipotypes considered.

In this study, we have focused exclusively on 5 cell types. Considering that several cell subtypes have recently been described, from different states of microglia to different cell subtypes based on the expressed proteins^10^, it is expected that in the case of lipotypes we will observe a diverse range not only based on these subtypes, but also considering the different states in which the cells are in compared to phenotypes, transcriptome or conectome. Therefore, this methodology demonstrates the potential to revolutionise our understanding of cellular diversity and function in the brain and, potentially, in other tissues, allowing the precise localisation not only of different lipotypes but also of distinct cell types without the need for specific protein markers, offering a novel approach to studying cellular composition. However, the current focus on a limited number of cell types and states suggests a potential limitation in capturing the full complexity of cellular heterogeneity, highlighting the need for further refinement and expansion of the lipotype library to encompass a wider range of cellular subtypes and states.

In conclusion, this study provides a universal tool to accurately determine the anatomical distribution of cell lipotypes, thus deepening our understanding of both the significance of the lipotypes themselves and the functions they may serve not only in the brain but in any tissue in health and disease.

## Methods

### Tissue samples

We used two-month-old male Sprague-Dawley rats (200–250 g) for the study. The animals were housed four or five per cage (50 cm length x 25 cm width x 15 cm height) at a temperature of 22°C and in a humidity-controlled (65%) room with a 12:12 hours light/dark cycle, with access to food and water ad libitum. This type of rats were used for immunohistochemical procedures and cell cultures analysed by MALDI mass spectrometry assays. Rats from different ages were used for primary cultures of the different cerebral cell types, including glial cells, neurons and choroid plexus. Thus, E18 Sprague-Dawley rat embryos were used for primary cultures of neurons, and newborn (P0-P2) Sprague-Dawley rats were used for cell cultures of astrocytes and microglia. The cell cultures of oligodendrocytes were obtained from dissected brains of P12 Sprague-Dawley rats. Finally, the choroid plexuses were obtained from 2-month-old rats. Moreover, to perform the lipotype-based image reconstruction in the rat model of basal forebrain cholinergic degeneration, we injected 192IgG-saporin (130 ng/µl) into the nucleus basalis magnocellularis (nbM), as previously described^5^. Control rats received an injection of artificial cerebrospinal fluid (aCSF) into the nbM. Rats were allowed to recover from surgery for 7 days. Two-month-old male C57BL/6 mice (25–35 g) were also used for mass spectrometry imaging, which were subsequently used for lipotype-based image reconstruction. The animals were housed four or five per cage (29 cm length x 16 cm width x 12 cm height) at a temperature of 22°C and in a humidity-controlled (65%) room with a 12:12 hours light/dark cycle, with access to food and water ad libitum. Every effort was made to minimise animal suffering and to use the minimum number of animals. All procedures were carried out in accordance with European animal research laws (Directive 2010/63/EU) and the Spanish National protocols that were approved by the Local Ethical Committee for Animal Research of the University of the Basque Country (CEEA 388/2014, CEEA M20-2018-52/54).

*Postmortem* human brain samples and data from donors included in this study were provided by Biobank of the Basque Country (CEISH/244MR/2015/RODRIGUEZ PUERTAS). The samples were obtained at autopsy after getting informed consent in accordance to the ethics committees of the University of Basque Country (UPV/EHU), following the Code of Ethics of the World Medical Association (Declaration of Helsinki), and guaranteeing the privacy rights of the human subjects. Specifically, samples of cortex and hippocampus from subjects with no previous pathology were used. These samples were also used for mass spectrometry imaging, which were subsequently used for lipotype-based image reconstruction.

### Primary brain cell cultures for the determination of the lipid composition (cell “lipotype”)

Cortical neurons were isolated from the cortical lobes of E18 Sprague-Dawley rat embryos, as described previously^11^, and resuspended in B27 Neurobasal medium plus 10% fetal bovine serum (FBS). Subsequently, neurons were seeded onto poly-L-ornithine coated glass coverslips at 10,000 cells/cm^2^. After 24h, the medium was replaced with serum-free (B27-supplemented Neurobasal® medium). The cultures were essentially free of astrocytes and microglia, and maintained at 37ºC and 5% of CO_2_.

Oligodendrocytes were isolated from optic nerves of P12 Sprague-Dawley rats, as previously described^12^, with minor modifications^13^. Cells were seeded onto poly-D-lysine coated glass coverslips at 10,000 cells per coverslip. The cultures were maintained at 37ºC and 5% CO_2_ in a chemically defined medium^14^.

Microglial cells were derived from cortical tissue of P0 - P2 Sprague-Dawley rats as previously described^15^. After 2 weeks, confluent monolayer of cultured astrocytes was removed from microglia by mechanical shaking. Free-floating microglial cells were purified by plating them on non-coated plastic Petri dishes (Sterilin™ plates). After 24h, non-adhered cells were eliminated and microglial cells were re-plated on poly-D-lysine coated coverslips.

Following these conditions, the purity of cultured microglia was higher than 99%. The cultures were maintained at 37ºC and 5% CO_2_.

Astrocytes were isolated from cerebral cortical tissue of P0–P2 Sprague-Dawley rats, as described elsewhere^16^. After 2 weeks, cells were trypsinised and astrocytes were plated onto poly-lysine cell culture vessels at 15,000 cells/cm^2^. The cultures were maintained at 37ºC and 5% CO_2_.

Choroid plexus cells were derived from choroid plexus epithelium. The tissues were trypsinised for 20 minutes and then centrifuged at 200g for 5 minutes. The supernatant was removed and the pellet was resuspended by mechanical digestion with Dulbecco’s Modified Eagle Medium (DMEM). Cells were seeded onto poly-D-lysine coated glass coverslips. The cultures were maintained at 37ºC and 5% CO_2_.

#### Preparation of cell homogenates for lipotype analysis

All cell cultures of different brain cell types were incubated with 500 µL of trypsin for 15 minutes with constant agitation. The supernatants were centrifuged at 16,000 rpm for 5 minutes at 4ºC. The pellets were resuspended in phosphate-buffered saline (PBS) and later were centrifuged at 16,000 rpm for 5 minutes at 4ºC. The resulting dry pellets were stored at -80ºC until the experiment was performed.

### Matrix - Assisted Laser Desorption Ionisation Mass Spectrometry Imaging (MALDI-MSI)

We used MALDI–MSI to describe the lipotype of the main different cell types present in the brain and to perform MSI of rat, mouse and human tissue. For the experiments aimed at identifying the lipotype of the cells, we used cell homogenates from the five main classes of cells present in the rat brain. For tissue MSI, fresh frozen tissue (from rat, mouse or human) was raised to -20°C, and 20 µm thick slices were cut in a cryostat (Microm HM550, Walldorf, Germany) and mounted onto gelatine-coated slides. These slices were kept at -25°C until used in MALDI-MSI assays. On the other hand, dry pellets from cell cultures (n=3-5) were thawed at 4ºC and resuspended in PBS. 5 - 10 µL of the suspensions were deposited onto gelatinised slides (n=14-40 dots per cell-type) and dried at room temperature to mimic the conditions of tissue sections measured by MALDI–MSI. Both types of samples, homogenates or sections, were dried in a glass sublimator (ACE glass 8023, USA) at room temperature for 15 min prior to deposition of the matrix onto the samples. Samples were sublimated with 300 mg of MBT in a sublimator at 100°C for 23 minutes, followed by a recrystallisation process on the bottom of a Petri plate with methanol at 38°C for one minute^17^.

MALDI–MSI analyses were carried out by using a LTQ–Orbitrap–XL mass spectrometer (Thermo Fisher Scientific, San Jose, CA, USA). The system was equipped with a nitrogen laser of λ = 337 nm, using a repetition rate = 60 Hz and a spot size= 80 × 120 μm. Each sample was scanned in positive ionisation mode, in the range of m/z 500 – 1000. The parameters were 2 μscans/step with 10 laser shots and a raster step size of 100-150 μm at laser fluence of 15 - 40 μJ.

A large number of molecules were detected, with some of them sharing very similar masses. This made it difficult to accurately assign an m/z value to a specific lipid molecule. Some of the m/z values obtained were assigned using different reference databases (e.g., Lipid Maps, www.lipidmaps.org or Human metabolome Database, www.hmdb.ca/). In addition, our data were also compared with previous publications from our research group and from others using similar methodological approaches. All the assignments were made with a mass accuracy of 5 ppm as the tolerance window.

### Immunohistochemistry

Immunohistochemistry assays were performed on fixed rat tissues. The 12 µm sections were simultaneously blocked and permeabilised with 4% donkey serum in 0.3% Triton X-100 in PBS (0.1 M, pH 7.4) for 2 h at room temperature (22 ± 2 °C). Sections were incubated overnight at 4 °C with antibodies that recognise specific proteins expressed by each cell type such as mouse anti-NeuN (1:250; Millipore) to label neurons, mouse anti-GFAP (1:500; Millipore) to label astrocytes and mouse anti-MBP (1:500; Biolegend) to label oligodendrocytes, all diluted in 0.3% Triton X-100 in PBS containing 5% BSA. The primary antibody was then revealed by incubation for 30 min at 37 °C in the dark with donkey anti-mouse Alexa Fluor-488 (1:250; Thermo Fisher Scientific) diluted in Triton X-100 (0.3%) in PBS.

### Lipotype-based image reconstruction

Analysis was performed using MATLAB with custom-made codes and freely available functions.

#### Cell-type spectrum

We first defined the representative spectrum of each cell-type as an average of the spectrum of the cells grown on purified cultures (see above tissues cultures).

For each slide scanned with MALDI and containing a given cell type colony, we first segmented the region of interests characterised by high cell density (HD-ROIs). For each pixel *(x,y)* within the HD-ROIs, the spectrum of peaks *SP(x,y,mz)*, where *mz* is the scanned mass, was extracted (using the library “imzMLConverter” compatible for MATLAB available at https://github.com/AlanRace/imzMLConverter; Reference AM Race, IB Styles, J Bunch, Journal of Proteomics, 75(16):5111-5112, 2012. doi.org/10.1016/j.jprot.2012.05.035) and normalised according to:

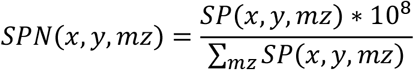

where SPN is the normalised spectrum.

The scanned mass range acquired across all samples was [500.011 1001.515] and, given the accuracy of the MALDI, it was binned with an interval of 0.012 *(bmz)* so for each pixel we computed a binned normalised spectrum (*b-SPN(x,y,bmz)*) where we summed the values of the *SP*(*x,y,mz*) following in the mass interval [*bmz-0*.*006 bmz+0*.*006*]. *NAN* (i.e. Not-A-Number) was assigned to *bmz* where no spectral value was acquired. Given a slide of a given cell type, an average spectrum was first calculated across all pixels, and finally a cell-type average spectrum (*AvS(bmz)*) was calculated as average across slides (blue plots in Extended Data Fig. 2a). Note that *NAN* were discarded, so averages included only numerical values. Extended Data Fig. 1 show an example for two slides of microglia of the *HD-ROIs*, and average *b-SPN* within the *HD-ROIs* and outside of them.

#### Definition of cell-type spectral signatures

In this work we considered five cell-types and each cell type was represented by its *AvS*_*ct*_*(bmz)*, where *ct* defines the specific cell type and is a scalar between 1 and 5 (cell-type order is: NEU, AS, OLI, MG and PLE, respectively). Since *AvSs* shared common spectral features as shown by the blues scatterplot in Extended Data Fig. 2b and by their high similarity (supra-diagonal part of the matrix shown in Extended Data Fig. 2c), we used a general linear model to quantify and remove spectral features shared across cell-types to extract a spectral signature for each cell type (*SpS*_*ct*_*(bmz)*). The general linear model only considered *bmz* for which a numerical value was present in all the *AvSs*, which represented about 9% of the mass bins. To calculate the *SpS*_*i*_*(bmz)* of a given cell type *i*, we discarded lower spectral values, so we considered only values of *AvSs* larger than 5*10^4^ (consider that major peaks were in the order of 10^6^), and we refer to this as *thresholded-AvS (T-AvS)*. The residuals (*R*_*i*_*(bmz)*) of the linear model estimating the T-*AvS*_*i*_*(bmz)* as the weighted combination of the other cell-types spectrum *T-AvS*_*j*_*(bmz)*, where *j*≠*i*, was first calculated. In correspondence of *bmz* where *R*_*i*_*(bmz)* was negative or *T-Asi(bmz)* was zero, *R*_*i*_*(bmz)* was set to zero and as a result *SpS*_*i*_*(bmz)* was obtained. These last two steps of processing the *R*_*i*_*(bmz)* were aimed to discard meaningless negative spectral values in cell-type spectral signature, and to remove spectral values which were previously thresholded in the AvS (being spectral values with low intensity).

Differently than the case of the *AvSs, SpS* display much less common spectral features with lower similarity across cell types (red scatterplots in Extended Data Fig. 2b and infra-diagonal part of the matrix shown in Extended Data Fig. 2c). Similarity shown in the matrix Extended Data Fig. 2c is calculated as cosine similarity^18^. SpSs and AvSs keep being highly correlated for each given cell type, as shown in the diagonal scatterplots and diagonal of the matrix of Extended Data Fig. 2b and c respectively.

#### Reconstruction of cell type distribution in brain tissue

The mass spectrum acquired in a specific point of a brain tissue *BTS(x,y,bmz)* was obtained first by normalising the spectrum acquired in a coordinate, i.e. pixel, *(x,y)* of the sample according to equation (1), then by binning in mass intervals of size 0.012 as explained above in the “Cell-type spectrum” section, and finally by thresholding with a value of 5*10^4^.

We used a linear model to estimate *BTS(x,y,bmz)* as linear combination of *SpS*_*ct*_*(bmz)* according to:

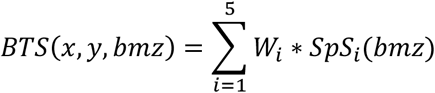

Only cell-types with positive weight contributions (for which *W*_*i*_>0) were considered for the cell-type presence estimation, and their contribution (C_i_) to the *BTS(x,y,bmz)* was obtained by normalising *W*_*i*_ to the sum of the positive weights, and multiplied by 100 to obtain a final percentage value according to:

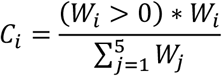

The estimation of overall similarity between *BTS(x,y,bmz)* and the linear combination of *SpS*_*ct*_*(bmz)* with positive weights defined as:

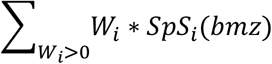

was computed using cosine similarity multiplied by 100 to obtain a percentage value.

## Data availability

All data associated with this study will be available to any researcher for the purpose of reproducing or extending the analysis. Initially, database from MALDI-MSI information will be generated and made accessible.

## Code availability

Code availability will be considered upon request in accordance with the legal terms of the participating institutions.

## Acknowledgements

Technical and human support provided by University of the Basque Country (UPV/EHU), Ministry of Economy and Competitiveness (MINECO), Basque Government (GV/EJ), European Regional Development Fund (ERDF), and European Social Fund (ESF) is gratefully acknowledged. J.M.-G. is the recipient of Margarita Salas fellowship funded by the European Union-Next GenerationEU. I.B.d.T is the recipient of Investigo fellowship funded by the European Union-NextGenerationEU. Cell-type illustrations were created with BioRender.com.

## Author information

## Authors Contribution

J.M.-G. and R.R.-P. conceived and designed the study, and together with P.B. performed the statistical analysis, and wrote the manuscript. C.M., A.L.-O and L.L. contributed to obtain cell cultures. J.M.-G., M.M.-R. and E.G.d.S.R. contributed to the performance of MALDI-MSI experimental procedures and data analysis. M.M.-R, I.M. and I.B.d.T. performed the immunofluorescences studies. J.S and P.B. contributed to computational development. All authors contributed to the manuscript revision, read, and approved the submitted version.

## Corresponding authors

J.M.-G., P.B. and R.R.-P

## Correspondence to

R.R.-P

## Ethics declarations

We agree with the statement of ethical standards required for manuscript submission. All procedures were performed in accordance with European animal research laws (Directive 2010/63/EU) and the Spanish National Guidelines for Animal Experimentation (RD 53/2013, Law 32/2007). The authors declare no conflict of interest and have approved the final article.

**Extended Data Fig 1.**
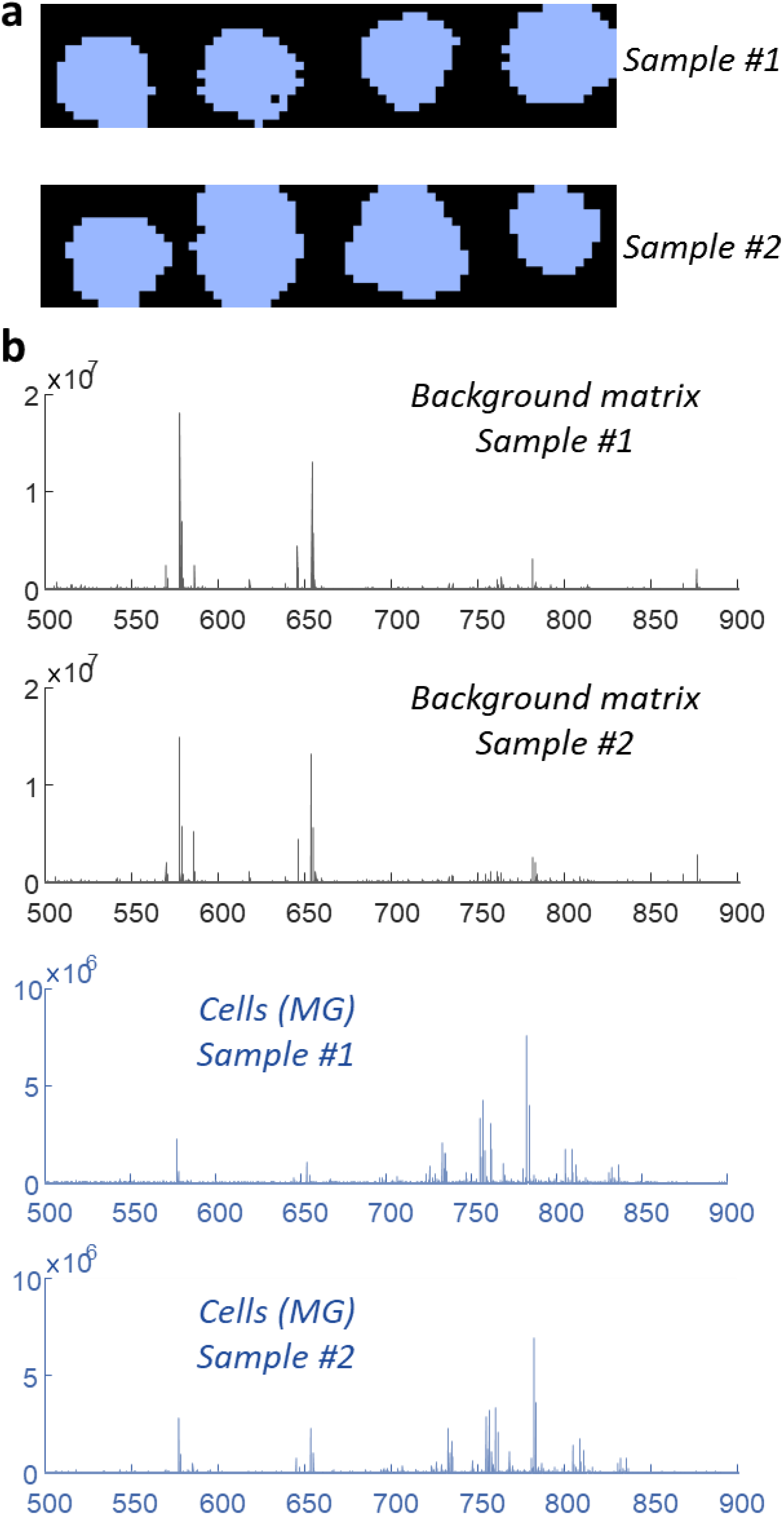
Extraction of spectrum from two different representative microglia cells. **a**, Two different representative images of the microglia pellets deposited on the slide are shown. The segmented blue area corresponds to dots with dense cell presence, from which the representative mass spectrum of the cell type is extracted (see blue plots in b). The black area corresponds to the absence of cells, which means that the acquired spectrum is related only to the coating molecular matrix (see black plots in b). **b**, Average mass spectrum extracted from the segmented background (top two black plots) and cell-type area (bottom two blue plots). The average is obtained across scanned pixels.

**Extended Data Fig 2.**
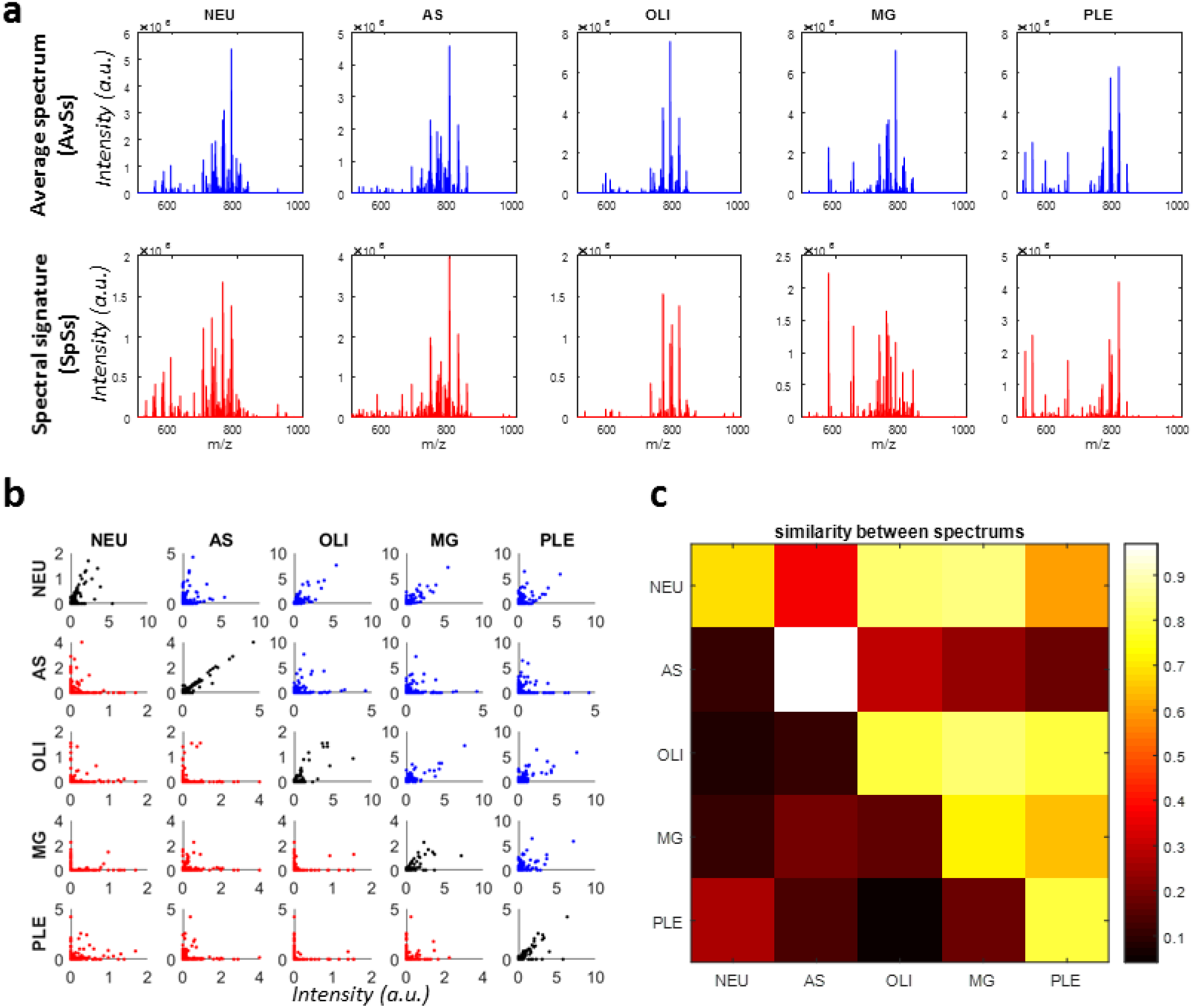
Construction of representative spectrum for each cell-type. **a**, In the top blue plots, the mass spectrum obtained as average through several cell cultures (AvS_ct_) for each cell type is shown. In the bottom red plots the spectral signatures (SpS_ct_) obtained through a linear model removing common spectral contribution are shown. Note that spectra are similar to the original ones although as shown in b and c, they lose their correlation across cell-types. **b**, For each pair of cell types (indicated on the top and left of the plots), a scatterplots of their spectral values for each common measured mass is shown. Blue scatterplots (supra-diagonal plots) are based on the AvS_ct_ while red scatterplots (infra-diagonal plots) are based on the SpS_ct_. Black scatterplots (diagonal plots) plot for each correspondent cell-type the SpS_ct_ (y-axis) vs the AvS_ct_ (x-axis).

**Extended Data Fig 3.**
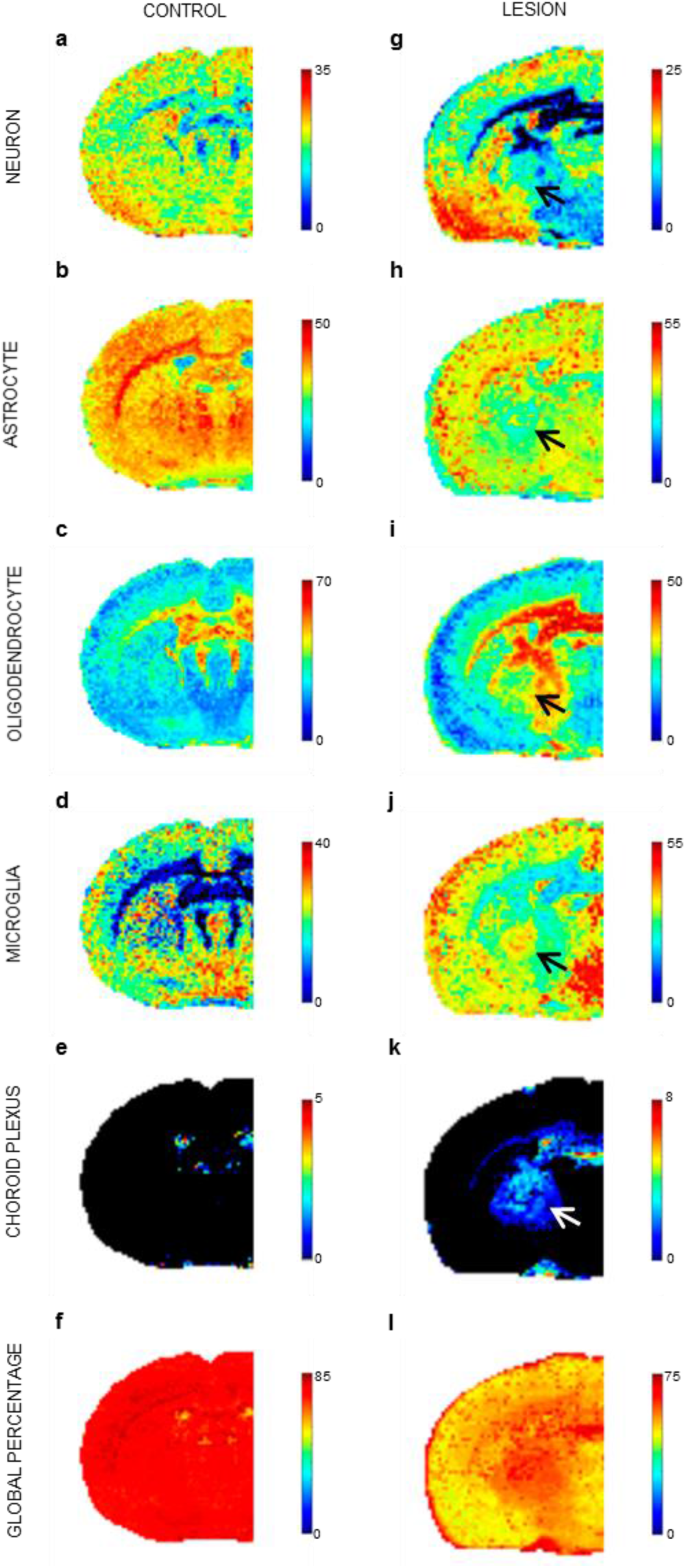
Reconstructed images of the localisation of the different lipotypes in coronal control (a-f) and toxin-induced lesion (g-l) rat brain section. Scale bar = 4mm. Note that the arrow shows how the lipotype image reconstructions show an increase in microglia (**j**) and cells present in the choroid plexus (**k**) at the lesion site, conversely, a decrease in astrocytes (**h**) is observed in the lesion area. It is noteworthy to highlight the observed changes in the image that represent the global percentage, suggesting potential changes in lipid composition within specific cell types, indicative of an activated state, e.g. in the case of choroid plexus cells, these changes could indicate cellular activation.

